# Evolutionary ecology of microbial populations inhabiting deep sea sediments associated with cold seeps

**DOI:** 10.1101/2022.05.09.490922

**Authors:** Xiyang Dong, Yongyi Peng, Muhua Wang, Laura Woods, Wenxue Wu, Yong Wang, Xi Xiao, Jiwei Li, Kuntong Jia, Chris Greening, Zongze Shao, Casey R.J. Hubert

## Abstract

Deep sea cold seep sediments host abundant and diverse bacterial and archaeal populations that significantly influence biogeochemical cycles. While numerous studies have revealed the community structure and functional capabilities of cold seep microbiomes, little is known about their genetic heterogeneity within species. Here, we examined intraspecies diversity patterns of 39 abundant species identified in sediment layers down to 4.3 mbsf across six cold seep sites from around the world. These species were predicted to participate in methane oxidation and sulfate reduction, and based on their metabolic capabilities, grouped as aerobic methane-oxidizing bacteria (MOB), anaerobic methanotrophic archaea (ANME) and sulfate-reducing bacteria (SRB). These physiologically and phylogenetically diverse MOB, ANME and SRB display different degrees of intrapopulation sequence divergence and different evolutionary trajectories. Populations were in general characterized by low rates of homologous recombination and strong purifying selection with most of the nucleotide variation being synonymous. Functional genes related to methane (*pmoA* and *mcrA*) and sulfate (*dsrA*) metabolisms were found to be under strong purifying selection in the vast majority of species investigated, although examples of active positive selection were also observed. These genes differed in evolutionary trajectories across phylogenetic clades but are functionally conserved across cold seep sites. Intrapopulation diversification of MOB, ANME and SRB species as well as their *mcrA* and *dsrA* genes was observed to be depth-dependent and undergo divergent selection pressures throughout the sediment column. These results highlight the role of the interplay between ecological processes and the evolution of key bacteria and archaea in deep sea cold seep sediments and shed light on how microbial populations adapt in the subseafloor biosphere.

## Introduction

Cold seeps are widely distributed along continental margins across the globe and likely have an important environmental influence on the local biology, chemistry, and geology^1–4^. These seep fluids are enriched in hydrocarbon gases (mainly methane) and other energy rich hydrocarbon fluids, and markedly alter the sedimentary microbial community structure and function by promoting microbial growth, specialization, and adaptation^1^. Based on amplicon sequencing of genetic markers and genome-resolved metagenomics, numerous studies have revealed the extensive macrodiversity (i.e. the measure of population diversity within a community^5^) of archaeal and bacterial lineages and microbial metabolic versatility along different cold seep sites and sediment depths^1, 6–9^. Findings include the discovery of various lineages of archaeal anaerobic methanotrophs (ANME) and sulfate-reducing bacteria (SRB) as syntrophic aggregates to perform methane oxidation coupled to sulfate reduction at anoxic sediment layers^10, 11^. In the upper oxic sediment layers of cold seeps, methane is mostly consumed by aerobic methane-oxidizing bacteria (MOB) mainly from the order *Methylococcales*^9, 12^. Although macro-variations in microbial diversity and functions have been well-characterized, our knowledge of the microdiversity (i.e. the measure of genetic variation within a population^13–15^) of these metabolically and taxonomically diverse subseafloor microorganisms remains limited. By addressing crucial questions such as “what populations have high microdiversity levels”, “which genes are under selection” and “does homologous recombination occur”, intra-population microdiversity analyses can provide a more complete understanding of microbial ecology and evolution in the subseafloor biosphere^16, 17^.

Traditional cultivation-based approaches have a fundamental role in studying genetic variation in microbial populations but are often not applicable to the study of the subseafloor biosphere as most microorganisms are extremely difficult to isolate^14, 18, 19^. With the emergence of bioinformatic tools for culture-free, high-resolution strain and subspecies analyses in complex environments^5, 15, 20–22^, genome-resolved metagenomic analyses can now be conducted at large-scale to reveal fine-scale evolutionary mechanism dynamics and strain-level metabolic variation^13, 14, 16, 17, 23–25^. Pioneering studies have been conducted in which metagenomic data has been used to explore the roles of basic processes (natural selection, mutation, genetic drift, and recombination) in shaping microbial evolution of several typical subseafloor habitats. For instance, *in situ* work examining genomic variation of microbes inhabiting the upper two meters of anoxic subseafloor sediments in Aarhus Bay revealed that rates of genomic diversification and selection do not change with either sediment age or depth, likely due to energy limitation and reduced growth in this environment^26^. In contrast to non-vent sediments, population-specific differences in selection pressure were observed in both *Sulfurovum* and *Methanothermococcus* species between two distinct geochemically distinct hydrothermal vent fields, where energy availability and cell abundances are relatively high^27, 28^. A third study found that gene flow and recombination appeared to shape the evolution of microbial metapopulations that disperse frequently through the cold, oxic crustal fluids of the mid-Atlantic ridge^29^. In contrast, no direct studies of the evolutionary histories and selection pressures of cold seep sedimentary microorganisms have been conducted despite the importance of their roles on global biogeochemical cycles. Knowledge of the nucleotide variation of key functional genes related to methane metabolism in these organisms is similarly lacking.

Here we hypothesize that microbial evolution in cold seep sediments may differ from observations in highly productive deep-sea hydrothermal vents and energy-limited marine sediments, due to the impact of the continuous flow of hydrocarbon-rich fluids. To gain insights into evolutionary trajectories among microbial populations inhabiting cold seep sediments, we examined the metagenomic data of 68 cold seep sediment samples to track population microdiversity from metagenomic short-read alignments and performed microdiversity-aware genomic comparisons. Our study also revealed the depth- and site-dependent trends of microbial evolution in deep sea cold seep sediments through the analysis of inter-sample genomic variation of species. Microbial adaptation in a gradient environment is a highly dynamic and complex process involving the interaction of multiple evolutionary forces^22^. Thus, the exploration of adaptive fingerprints to uncover evolutionary mechanisms of specific taxa in cold seep sediments may hint at the factors impacting long-term evolution in deep subseafloor biosphere, as well as those processes that shape the evolution of genes involved in adaptation to specific environmental factors.

## Results and Discussion

### Depth profiles of species-level clusters in cold seep sediments

We assembled metagenomic data sequenced from 68 sediment samples obtained from six globally distributed cold seep sites, spanning various depth layers from 0 to 4.3 mbsf (**Supplementary Figure 1 and Supplementary Table 1**). After binning of metagenomic assemblies, 1261 prokaryotic species-level clusters (1041 bacterial and 220 archaeal; **Supplementary Table 2**) were recovered according to the suggested threshold of 95% average nucleotide identity (ANI) for delineating species^30–32^. These species clusters belonged to 85 phyla (70 bacterial and 15 archaeal; **Figure 1a**) based on the Genome Database Taxonomy (GTDB; version R06-202)^33–36^, and were highly represented by bacterial phyla including Chloroflexota (n = 184), Proteobacteria (n = 125), Desulfobacterota (n = 101), Planctomycetota (n = 73) and Bacteroidota (n =67). The top five archaeal phyla with the largest number of species-level clusters were Asgardarchaeota (n = 50), Thermoplasmatota (n = 44), Halobacteriota (n = 42), Thermoproteota (n = 35; mainly *Bathyarchaeia*) and Nanoarchaeota (n = 19). Most of these phyla lack an available cultured representative in the GTDB^35^ and consist exclusively of MAGs and/or SAGs. Around 51% and 94% clusters could not be assigned to an existing genus and species, respectively (**Figure 1b**), confirming that the majority of the cold seep sediment species lack even uncultured representative genomes in the reference database.

**Figure 1.**
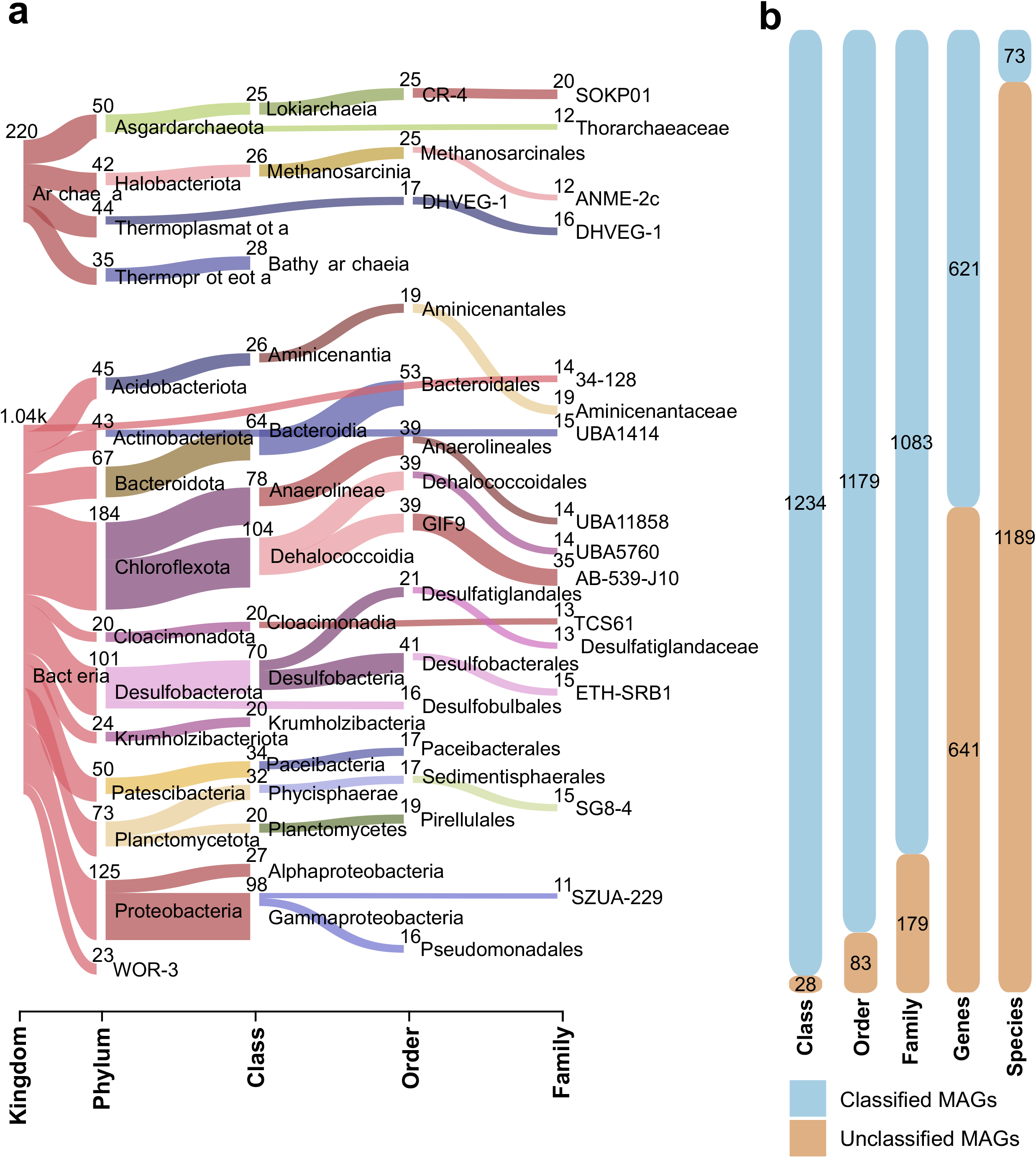
Classification of species-level MAGs recovered from global cold seep sediments. (a) Sankey based on assigned GTDB taxonomy showing recovered archaeal and bacterial MAGs at different phylogenetic levels. Numbers indicate the number of MAGs recovered for the lineage. (b) Total MAGs unclassified by GTDB-Tk at each taxonomic level. MAGs were dereplicated at species level (i.e., 95% ANI). Detailed statistics for 1261 MAGs are provided in **Supplementary Table 2**.

At the phylum level, the total relative abundance of Halobacteriota was the highest across samples of all eight depth groups (**Supplementary Figure 2**), and ranged from 2.9%±3.7% at 0-0.05 mbsf (n = 11) to 20.7%±12.6% at 2.0-3.0 mbsf (n = 4) (**Supplementary Table 4**). At the species level (**Supplementary Figure 3**), clusters from ANME-1 (genus of QEXZ01) ranked the highest (up to 9.1%±9.1% at 0.3-4.5 mbsf), followed by ANME-2c groups (up to 2.0%±2.3% at 0-0.3 mbsf), indicating their different depth distributions^37^. Bacterial taxa belonging to the phyla Desulfobacterota and Caldatribacteriota are also very abundant, with the latter being especially prevalent in deeper sediments (**Supplementary Figure 2**). The most abundant bacterial species was assigned to Caldatribacteriota taxon SB_S5_bin2, with values of up to 2.2%±2.3% at 1.0-2.0 mbsf (**Supplementary Figure 4**). Three species belonging to Desulfobacterota, the potential bacterial partners of ANME-1 or ANME-2^38^, became relatively predominant (up to 3.5%±0.5%) at 2.0-4.5 mbsf. Overall, sediment depth is one important factor shaping distributions of cold seep sediment microbial species clusters^7, 39^.

### Selection of key functional taxa for microdiversity analysis

Aerobic methane-oxidizing bacteria (MOB), anaerobic methanotrophic (ANME) archaea, and sulfate-reducing bacteria (SRB) are three key groups of functional microorganisms in cold seeps sediments^1, 6, 9^. To identify them, we screened the 1261 MAGs for the presence of three functional genes: the particulate methane monooxygenase marker gene *pmoA* encoding for MOB, the methyl-coenzyme-M reductase marker gene *mcrA* for ANME archaea, and the dissimilatory sulfite reductase marker gene *dsrA* for SRB. To facilitate the microdiversity analyses of key functional taxa, we kept only species-cluster representative MAGs with an estimated quality score ≥50 (defined as the estimated completeness of a genome minus five times its estimated contamination)^40^ and at least 10× coverage^15, 22, 41, 42^.

Three species belonging to *Gammaproteobacteria* were found to contain *pmoA* genes and passed the required criteria for MAG quality (**Figure 2a and Supplementary Table 3**). They were found exclusively in near-surface sediments (0-0.1 mbsf) from the Haima cold seep in the South China Sea (**Supplementary Figure 5 and Supplementary Table 5**), where oxygen is likely still available via penetration from the water column. Functional annotation of 13 species genomes from *Methanosarcinia* and *Syntropharchaeia* indicated the ability of these species to perform anaerobic oxidation of methane (**Figure 2b and Supplementary Table 3**). These 13 species represent four families: *Methanosarcinaceae,* HR1, ANME-2c and ANME-1, and are widely distributed in 53 samples across sediment depths in the range of 0.01-4.25 mbsf (**Supplementary Figure 5 and Supplementary Table 5**). Among 117 *dsrA*-containing species genomes, 23 bacterial species from Desulfobacterota perform sulfate reduction, likely coupled to anaerobic oxidation of methane **(Figure 2c and Supplementary Table 3)**, and belonged to four clades: “C00003060” (aka SEEP-SRB1c^43^), *Desulfobacterales, Desulfobulbia* and *Desulfatiglandales*. They are widely distributed in 12 sediment columns and along multiple sediment depths (0.01-4.25 mbsf) at various relative abundances (**Supplementary Figure 5 and Supplementary Table 5**).

**Figure 2.**
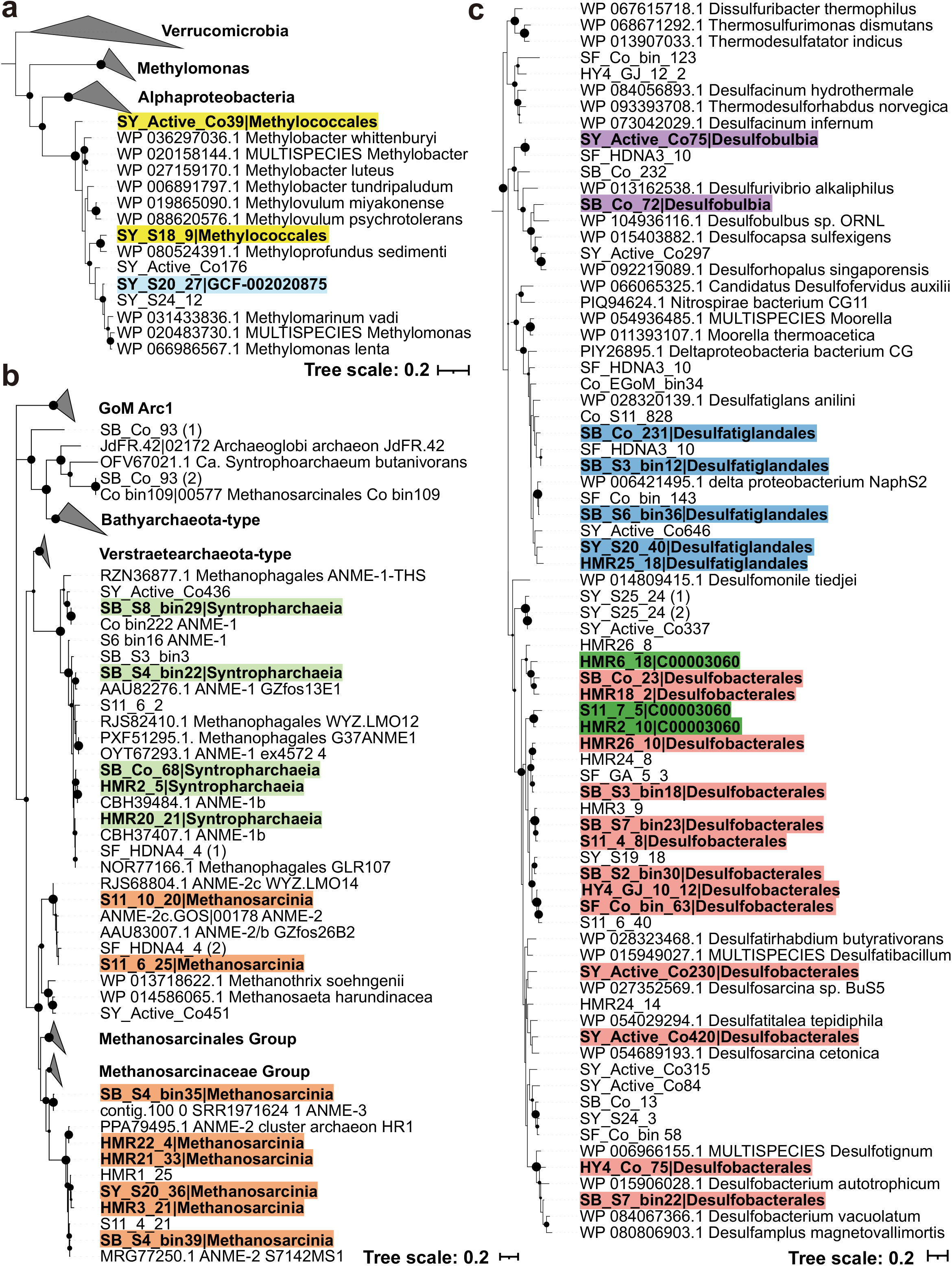
Maximum-likelihood phylogenetic trees of three key functional genes in cold seep sediments. Phylogenetic trees are based on alignments of amino-acid sequences of (a) PmoA, (b) McrA and (c) DsrA protein sequences. The sequences from the same taxonomic groups are highlighted in the same colors. Black dots indicate bootstrap values 50-100%. Scale bars indicate the average number of substitutions per site. Detailed annotations of species-cluster representative MAGs are provided in **Supplementary Table 3**.

### Genomic variations across different phylogenetic groups

At the genome level, we evaluated the 39 functional species for linkage disequilibrium (D’), pN/pS (the ratio of non-synonymous/synonymous mutations), r/m (the rate of recombination to mutation, gamma/mu; only for 10 species, **see Methods**), and number of single nucleotide variations for every thousand base pairs (SNVs/kbp). For populations from MOB, ANME and SRB, the evolutionary metrics varied greatly (**Figure 3a and Supplementary Table 5**), showing a wide distribution range in D’ (~0.89 vs 0.72-0.99 vs 0.74-0.99), SNVs/kbp (5-41 vs 2-98 vs 2-89) and pN/pS (0.12-0.27 vs 0.11-0.25 vs 0.13-0.23). D’ values indicate that MOB, ANME and SRB have not undergone high rates of homologous recombination similar to soil bacterial populations across a grassland meadow^17^. Population diversity in these groups as measured by SNVs/kbp is higher than that observed for soils in grassland meadows and subseafloor crustal fluids^17, 29^. These result indicate that deeply buried cold seep populations from MOB, ANME and SRB are under purifying selection, suggesting the possibility that these populations have reached an adaptive optimum for this stable environment, which is maintained by purging nonsynonymous mutations^44^. This result is in line with previous observations reported for bacterioplankton assemblages in sunlit freshwater and marine systems^16, 24, 45^, and microbial populations in other deep sea biospheres^46, 47^.

**Figure 3.**
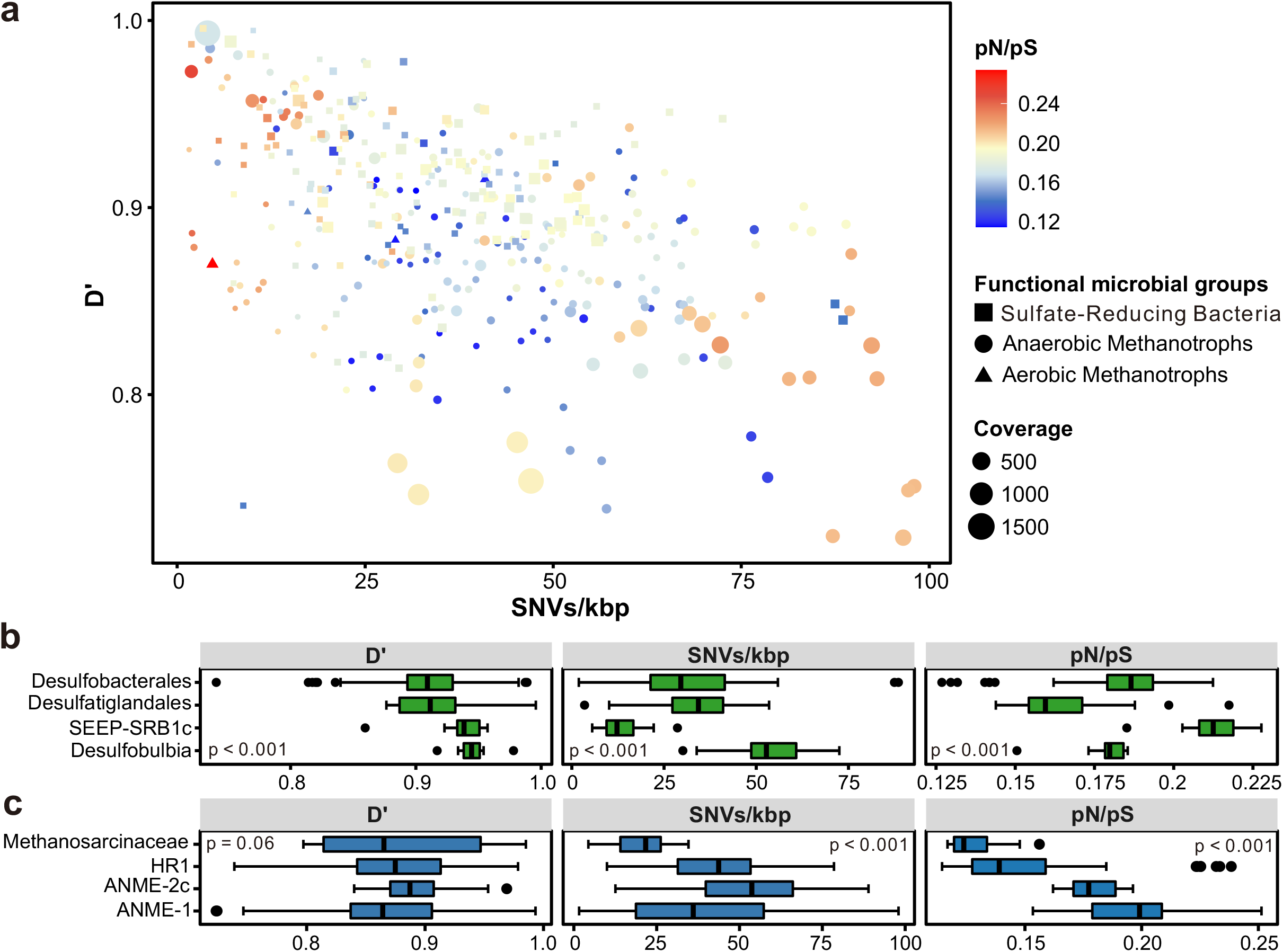
Genome-wide evolutionary metrics of three key functional microbial groups in cold seep sediments. (a) Relationships between SNV density (SNVs/kbp), linkage disequilibrium (D’), the ratio of nonsynonymous to synonymous polymorphisms (pN/pS ratio), and genome coverage at genome level. Each dot represents one species-level microbial population. (b)-(c) Box plots showing comparison of SNV density, D’ and pN/pS of sulfate-reducing bacteria and anaerobic methanotrophic archaea across different taxonomic groups. P-values of differences across different taxonomic groups were calculated using Kruskal-Wallis rank sum test. Source data is available in **Supplementary Table 5**.

These data (**Figure 3a and Supplementary Table 5**) also indicate that microbial populations with different functional features in cold seep sediment habitats have diverse evolutionary modes, similar to observations in deep-sea hydrothermal vents with unique attributes^13, 27^. Accordingly, statistically significant differences were observed for almost all of the evolutionary metrics among the ANME and SRB lineages (*P*<0.001, except 0.06 for D’ of ANME groups; **Figures 3b and 3c**). The SEEP-SRB1c group had the lowest SNVs/kbp ratio, highest pN/pS value and a low degree of recombination among the four SRB groups (**Figure 3b**). Fewer recombination events and lower nucleotide diversity are signals of selective sweep^24^. Thus, these data indicate that the SEEP-SRB1c group has undergone strong selection.

For HR1, *Methanosarcinaceae*, ANME-2c, *Desulfatiglandales* and ANME-1, negative correlations between D’ and SNVs/kbp reflect a positive relationship between nucleotide diversity and homologous recombination **(Figure 4a and Supplementary Table 6)**. This is in agreement with the positive correlation (linear regression; *R^2^*=0.34, *P*<0.001) found between r/m and SNVs/kbp for both ANME and SRB **(Supplementary Figure 6a and Supplementary Table 7)**. The ratio of nucleotide substitutions originating from homologous recombination to those originating from mutation (r/m ratio) can be used to measure the relative effect of homologous recombination on the genetic diversification of populations^48^. These results indicate that ANME and SRB populations could preserve high genome-wide diversity and prevent selective sweeps through increasing recombination rates to various environmental changes^14, 24^. In addition to the negative correlation between SNVs/kbp and D’, negative correlations were also observed between SNVs/kbp and pN/pS for HR1 and *Methanosarcinaceae* **(Figure 4b, Supplementary Figure 7 and Supplementary Table 6)**. These correlations indicate that HR1 and *Methanosarcinaceae* populations are stabilized by frequent recombination while maintaining a high degree of intra-population diversity characterized by an accumulation of synonymous mutations, pointing to an ancient divergence of these two populations^45, 49^. In contrast, the *Desulfobulbia* populations had higher pN/pS values with more single nucleotide variants (linear regression; *R^2^*=0.65, *P*=0.006; **Figure 4b and Supplementary Table 6)**. Additionally, they also had high SNVs/kbp and low degrees of within-species recombination **(Figure 3c and Supplementary Figure 8)**. These data suggest that *Desulfobulbia* populations may be in the process of subspecies establishment (i.e. speciation) or purging of non-synonymous mutations^14^. For ANME-2c and *Desulfobacterales* populations, the genome coverage (i.e. relative abundances of the populations) and SNVs/kbp fitted the linear regression model with a positive slope **(Figure 4c and Supplementary Table 6)**, indicating that population quantity may be constraining genomic microdiversification^42^. ANME-2c populations with higher abundances were found to show high single-nucleotide variations which were related to the high mutation rate or accumulation of mutations in the population (**Supplementary Figure 7**)^27, 50^. Relatively constant pN/pS ratios in those population further suggest that non-synonymous mutations in ANME-2c population might have been purged by purifying selection over a long period^45^. For the *Desulfobacterales* group, high-coverage populations were also reported to show relatively high degrees of recombination, and recombination did not bring changes in amino acids at the genome level despite the observed high nucleotide variations (**Supplementary Figure 8**)^22^. ANME-1 populations with higher abundances showed higher recombination rates (linear regression; *R*^2^=0.12, *P*<0.001; **Figure 4d and Supplementary Table 6**), consistent with the positive linear regression relationship between r/m and coverage (linear regression; *R^2^*=0.81, *P*<0.001; **Supplementary Figure 6b and Supplementary Table 7**). These findings are consistent with those of another study in which the relationship between population abundance and recombination rate was proposed to be the underlying mechanism responsible for the evolutionary success of the marine bacterium SAR11 in the near-surface epipelagic waters of the ocean^45^.

**Figure 4.**
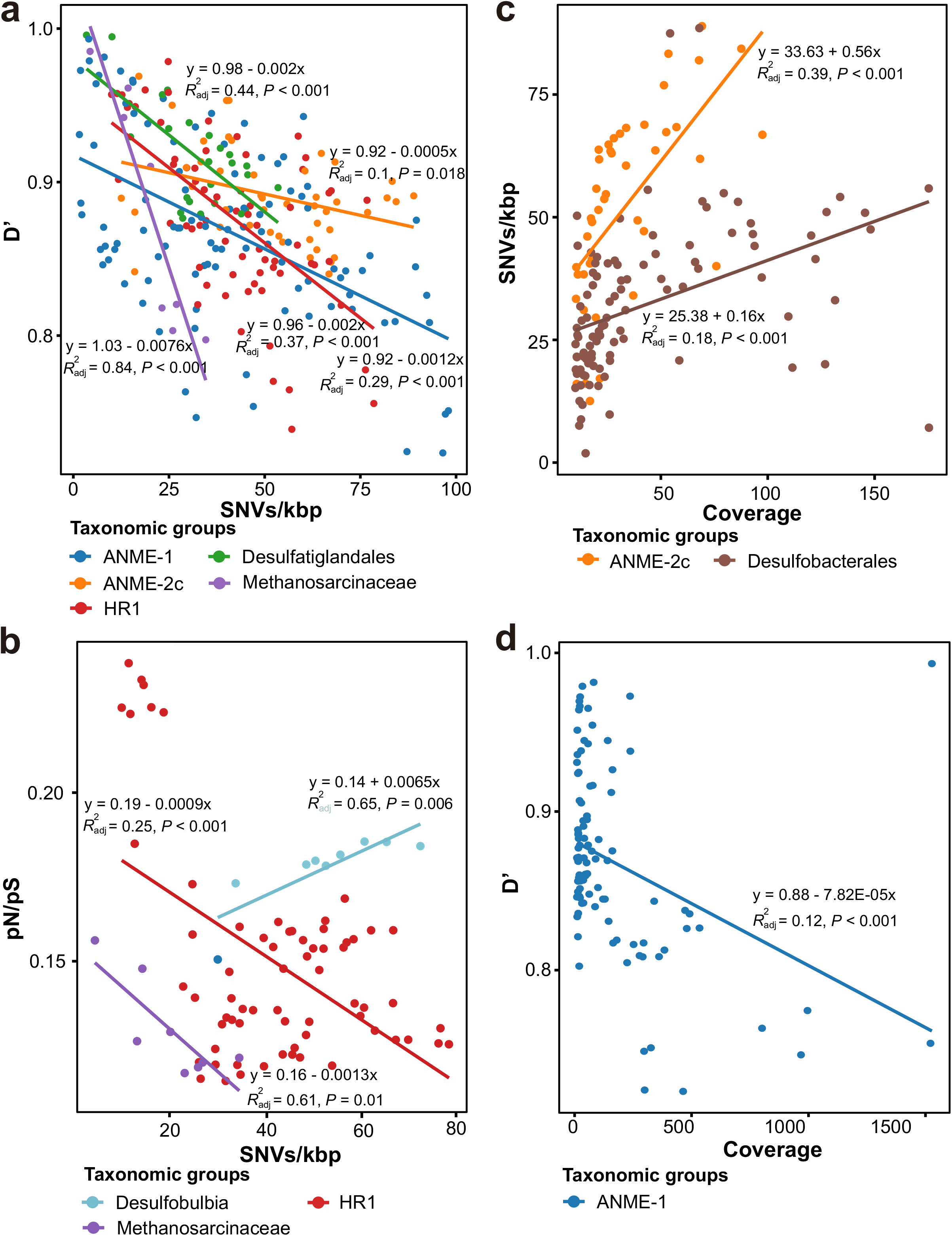
Genome-wide comparison of evolutionary metrics for microbial populations in cold seep sediments. (a)-(b) D’ and pN/pS ratio in relation to SNV density. (c)-(d) SNV density and D’ in relation to genome coverage. Each dot represents one species-level microbial population. Linear regressions and R^2^ values are indicated for different taxonomic groups. Detailed statistics for linear regressions are provided in **Supplementary Table 6**.

### Nucleotide variation of three key functional genes

For *pmoA, mcrA* and *dsrA* genes, SNVs/kbp ranged widely from 0.76 to 123 (45 on average, **Figure 5a and Supplementary Table 8)**. Based on the pN/pS values (0-1.43, 0.16 on average), these genes were under strong purifying selection. The evolutionary fitness of these three key functional genes was also consistent with that observed in previous studies for natural comammox *Nitrospira* populations^44^ and *Thalassospira* bacterial populations isolated from million-year-old subseafloor sediments^19^. Our findings are in line with research showing that essential genes and enzymes catalyzing reactions that are difficult to bypass through alternative pathways are subject to higher purifying selection than nonessential ones^26, 44^. Even though the pN/pS values of the vast majority were well below 1 (indicating purifying selection), genes with pN/pS values above 1 and significantly higher than the genomic average were detected, which indicates that positive selection acted upon those genes. Although these microorganisms were deeply buried in subseafloor in which microbial evolution might operate differently from sunlit habitats^13^, the distribution profile of pN/pS values are compatible with the neutral theory model^51^ wherein most mutation events are neutral or deleterious^52^ (**Figure 5b**). Similarly, in microbial rare genes, evolution was found to proceed largely via neutral processes^53^. On the other hand, studies of the microbial inhabitants of wild bromeliads demonstrate patterns indicating the action of non-neutral processes^54^.The similar evolutionary patterns appeared in both microbes and higher eukaryotes may be due to the conserved ancient mechanism.

**Figure 5.**
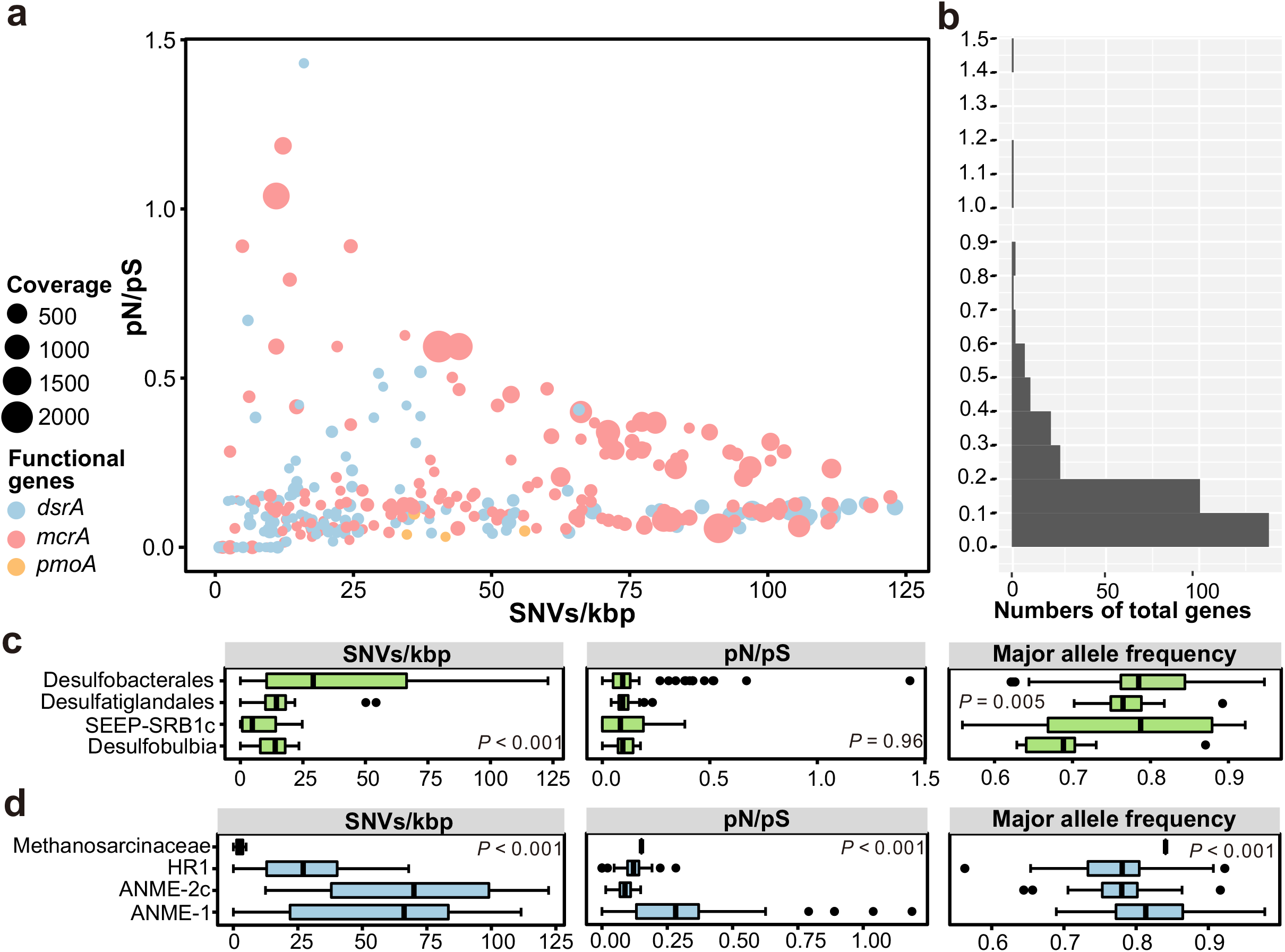
Gene-specific evolutionary metrics of three key functional microbial groups in cold seep sediments. (a) Relationships between SNV density, pN/pS and gene coverage at gene level. Each dot represents one species-level microbial population. (b) Frequency histogram of pN/pS. (c)-(d) Box plots showing comparison of SNV density, pN/pS and major allele frequency of *dsrA* and *mcrA* genes across different taxonomic groups. P-values of differences across different taxonomic groups were calculated using Kruskal-Wallis rank sum test. Source data is available in **Supplementary Table 8**.

For *dsrA* genes, SNVs/kbp and major allele frequency exhibited statistically significant differences among the four SRB groups (*P*≤0.005), while pN/pS values were similar among them **(Figure 5c)**. Similar evolutionary trends (relatively low SNVs/kbp and pN/pS values) were observed in *dsrA* genes from groups of *Desulfobulbia,* SEEP-SRB1c and *Desulfatiglandales,* **(Supplementary Figure 9)**. This indicates that *dsrA* genes in these groups were functionally stable following purifying selection. The *dsrA* gene in the *Desulfobacterales* group also had low pN/pS values (~0.12) but showed a broad range of SNVs/kbp. The *dsrA* gene of ETH-SRB1 SB_S7_bin23 populations with abnormally high pN/pS values were found with fewer synonymous mutations **(Supplementary Figure 10)**, indicating that *dsrA* genes in these populations were under strong positive selection and were likely further determined by random genetic drift^55^.

For *mcrA,* the three evolutionary metrics (SNVs/kbp, pN/pS and major alleles frequency) were significantly different (*P*<0.001) among the four ANME groups **(Figure 5d and Supplementary Figure 11)**, indicating differences in evolutionary trends of *mcrA* genes across these groups. The *mcrA* gene from HMR20_21 in the ANME-1 group were found to have high pN/pS values with low synonymous mutation rates, indicating positive selection or relaxed purifying selection **(Supplementary Figure 10)**. For *mcrA* genes from ANME-2c, SNVs/kbp (linear regression; *R^2^*=0.47, *P*<0.001) and pN/pS values (linear regression; *R^2^*=0.17, *P*=0.003) positively correlated with gene coverage **(Supplementary Figure 12)**. This suggests that mutations were maintained for *mcrA* genes during the clonal expansion of ANME-2c populations.

Major allele frequency (0.79 on average) appeared to show a direct relationship with SNVs/kbp for *mcrA* and *dsrA* genes (**Supplementary Figures 9 and 11**). High SNVs/kbp corresponded to high major allele frequencies (~0.8), while the distributions of major allele frequency were mostly scattered (0.56-0.99) at low values of SNVs/kbp. Specific major alleles were fixed in most *mcrA* and *dsrA* genes (major allele frequency: 0.70-0.98) with varying degrees of nucleotide diversity (SNVs/kbp: 1-123). For *mcrA* and *dsrA* genes with relatively low major allele frequency (0.56-0.70) and low pN/pS values (0-0.26), their genetic heterogeneity was preserved and strongly selected in populations from HR1, SEEP-SRB1c, *Desulfobulbia* and *Desulfobacterales* which might help diversification of functional genes under different environmental conditions^56^.

### Depth- and site-dependent trends of microdiversity

To determine whether microbial microdiversity was depth-dependent in deep-sea cold seep sediments, we assessed the relationship between evolutionary metrics and sediment depth. At genome level, a negative correlation was observed between SNVs/kbp and depth while pN/pS and D’ showed a positive relationship with sediment depth for ANME and SRB populations **(Figures 6a-c and Supplementary Table 9)**. This suggests that as depth below the sea floor increases along the sediment column, microbial populations exhibit less microdiversity and levels of homologous recombination, as well as more relaxed purifying selection. This depth-dependent trend differs from that found for non-seep subseafloor sediments in which buried microbial populations show uniformly low genetic heterogeneity across sediment depths^26^; but similar to results from a pelagic freshwater system on the surface of the Earth where a lower mutation rate was observed in the deeper water layer^42^. This is likely related to the higher energy supply and larger population sizes in cold seep sediments compared to non-seep subseafloor habitats^29^. Depth-dependent microdiversity patterns were also observed for *mcrA* and *dsrA* genes. In general, SNVs/kbp and pN/pS are negatively correlated with sediment depth while major allele frequency correlated positively with depth but the degree to which it did so was population-specific **(Figure 6d-f and Supplementary Table 9)**. This suggests that *mcrA* and *dsrA* genes had lower degrees of microdiversity and were subject to higher levels of purifying selection when ANME and SRB were buried deeper. Additionally, based on observations at genome and gene levels, ANME and SRB populations likely undergo distinct selection pressures arising from sediment depths **(Figure 6)**.

**Figure 6.**
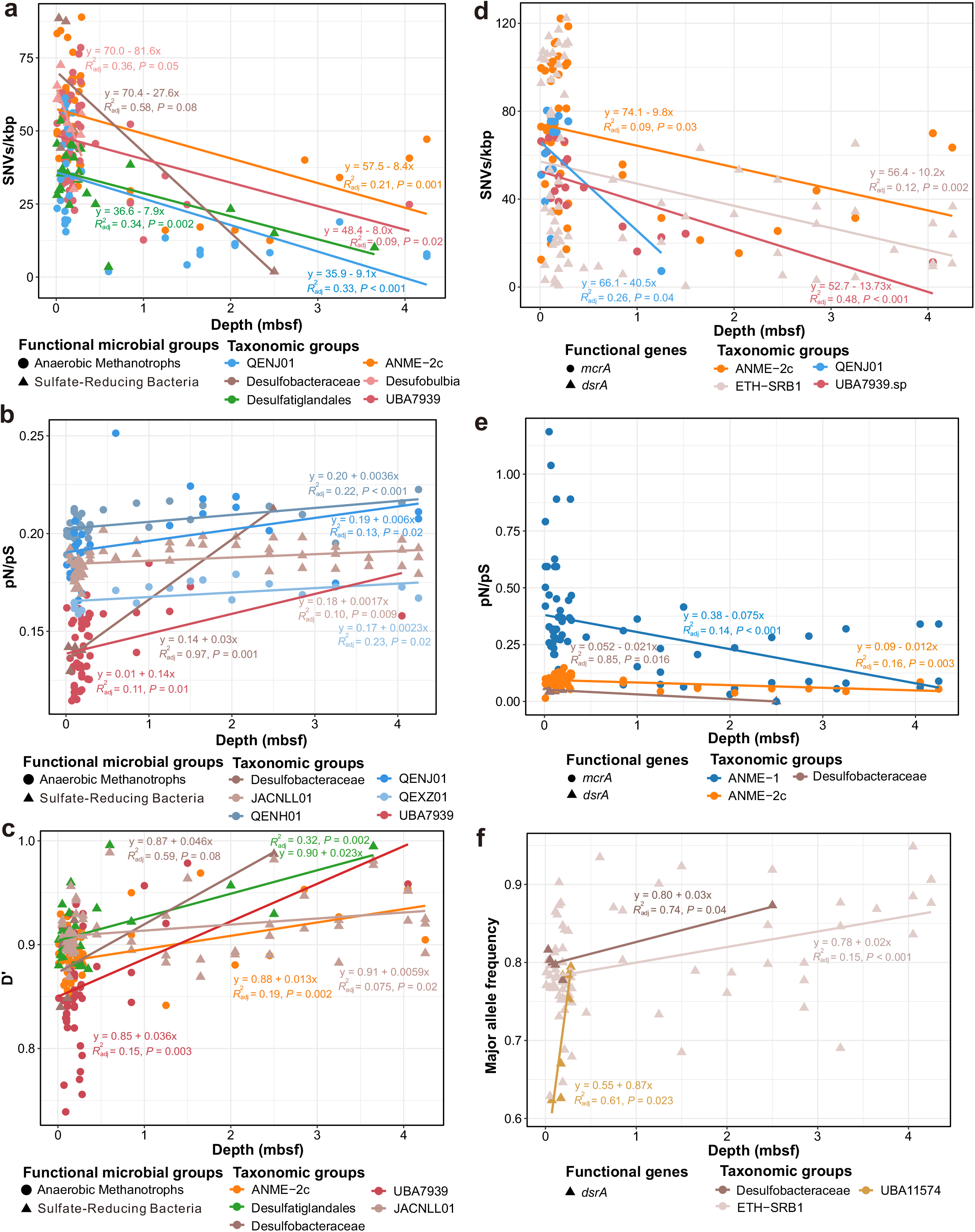
Relationships between evolutionary metrics and cold seep sediment depths (mbsf) at the whole-genome and gene levels. (a)-(c) Comparison of SNVs/kbp, pN/pS ratio and D’ against sediment depths at the whole-genome level. (d)-(f) Comparison of SNVs/kbp, pN/pS ratio and major allele frequency against sediment depths at the gene level. Each dot represents one species-level microbial population. Linear regressions and R^2^ values are indicated for different taxonomic groups. Detailed statistics for linear regressions are provided in **Supplementary Table 9**.

Cold seep microbiomes are reported to be locally selected and diversified (macrodiversity) by unique benthic biogeochemical conditions and environmental gradients such as for methane and sulfate concentrations^6, 7, 9, 39^. Evolutionary metrics (D’, SNVs/kbp, pN/pS) at the genome level showed significant differences (*P*<0.001) among different cold seep sites **(Supplementary Figure 13a)**, indicating that different physicochemical conditions of various cold seep systems also influenced the intra-population diversity and evolutionary processes in microbial populations. We also observed that there were differences (*P*=0.006) in evolutionary metrics (SNVs/kbp and major alleles frequency) for the three functional genes among different cold seep sites, **(Supplementary Figure 13b),**indicative of site-dependent microdiversity for functional genes. However, no clear difference (*P*=0.64) was observed among them with regard to pN/pS **(Supplementary Table 10)**. This suggests that those three key genes from ANME and SRB are mostly functionally conserved across different cold seep sites.

### Conclusion

By analyzing a suite of population genetics parameters through metagenomic read mapping, here we show that microbial evolutionary processes of microbes in deep-sea cold seep sediments are much more complex than previously thought, and are governed by factors that differ from those observed in energy-limited marine sediments, hydrothermal vents and low biomass subseafloor fluids^26, 27, 29^. The 39 abundant MOB, ANME and SRB species in cold seep sediments had diverse evolutionary modes with different degrees of nucleotide variation and varying degrees of homologous recombination, demonstrating that selection pressure exerted by seep fluids enriched in methane may operate differently on different species-level populations. The investigated species from MOB, ANME and SRB in general showed low homologous recombination and strong purifying selection, the latter process being especially strong in functional genes related to methane (*pmoA* and *mcrA*) and sulfate (*dsrA*) metabolisms. Evolutionary metrics of these genes differed across species identities but were functionally conserved across various cold seep sites, supporting that the importance of relatively stable cold seep environmental conditions (i.e. continuous supply of methane and sulfate) in affecting evolutionary processes of genes essential in energy metabolism. We further found that sediment depths can not only shape the community structure of cold seep microbes in sediment depth layers from 0 to 4.3 mbsf, but also be one of the driving factors of microdiversity patterns for microbial genomes and genes. The depth-dependent trends of microdiversity might be the mixing consequences of redox condition changes and age of the sediment^12, 26^. Together, this study improves our understanding of principles that drive evolution of slow-growing deep-sea microbes in one of the unique subseafloor biosphere. However, these conclusions were only based on abundant populations from MOB, ANME and SRB, genomic microdiversification of other species at lower abundance levels can still play important functional roles was not evaluated due to relatively shallow sequencing^5^. Deeper sequencing is therefore needed to depict a full picture of microdiversity within microbial populations in deep sea sediments^57^.

## Methods

### Metagenomes for deep-sea cold seep sediments

Metagenomic data sets were compiled from 68 sediment samples (0 to 4.3 mbsf) collected from six globally distributed cold seep sites (**Supplementary Figure 1**). These sites are as follows: Eastern Gulf of Mexico; Northwestern Gulf of Mexico; Scotian Basin; Haiyang4, Site F, and Haima cold seeps in the South China Sea (**Supplementary Table 1**). For samples from Northwestern Gulf of Mexico, metagenomic data sets along with metadata were downloaded from NCBI Sequencing Read Archive (SRA) and NCBI BioSample databases^58^. Samples of SY5, SY6 and S11 were obtained from sediments of Haima cold seep areas^59^ and raw sequencing data were deposited in NCBI SRA (PRJNA739036 and PRJNA738468). For other samples, sample collection and DNA sequencing were detailed previously^7, 8, 60–62^. The 68 sediment samples were catalogued into eight groups according to depth below the seafloor (**Supplementary Figure 2**): 0-0.05 mbsf (n = 11); 0.05-0.1 mbsf (n = 14); 0.1-0.2 mbsf (n = 16); 0.2-0.3 mbsf (n = 8); 0.3-1.0 mbsf (n = 6);1.0-2.0 mbsf (n = 4); 2.0-3.0 mbsf (n = 4); 3.0-4.5 mbsf (n = 5).

### Metagenome assembly and binning

Paired-end raw reads were quality-controlled by trimming primers and adaptors and filtering out artifacts and low-quality reads using the Read_QC module within the metaWRAP pipeline (v1.3.2; –skip-bmtagger)^63^. For each cold seep site, filtered reads from each metagenome were individually assembled and co-assembled using MEGAHIT (v1.1.3; default parameters) based on succinct *de Bruijn* graphs^64^. Contigs less than 1000 bp were removed. For each assembly, contigs were binned using the binning module (parameters: –maxbin2 –concoct –metabat2) and consolidated into a final bin set using the Bin_refinement module (parameters: –c 50 –x 10) within metaWRAP. The quality of the obtained MAGs was estimated by the lineage-specific workflow of CheckM (v1.0.12)^65^. MAGs estimated to be at least 50% complete and with less than 10% contamination were retained.

### Species-level clustering and taxonomic assignment

Species-level clustering and representative species identification was performed using dRep (v3.2.2)^31^ with an average nucleotide identity (ANI) cutoff value of 95%. The taxonomic classifications of representative MAGs were assigned based on the Genome Database Taxonomy GTDB (release 06-RS202)^35^ via the classify workflow of GTDB-Tk (v1.5.1)^66^. To calculate the relative abundance of each MAG, CoverM was used in genome mode (v0.6.0; parameters: –min-read-percent-identity 0.95 –min-read-aligned-percent 0.75 –trim-min 0.10 –trim-max 0.90; https://github.com/wwood/CoverM).

### Functional annotations and phylogenetic analysis

METABOLIC-G, an implementation of METABOLIC (v4.0)^67^, was used to predict metabolic and biogeochemical functional trait profiles of MAGs. For phylogenetic analysis of functional genes related to methane and sulfate metabolism, amino acid sequences were aligned using the MUSCLE algorithm^68^ included in the software package MEGA X^69^. All positions with less than 95% site coverage were excluded. The maximum-likelihood phylogenetic tree was constructed in MEGA X using the Jones Taylor Thornton matrix-based model, bootstrapped with 50 replicates. The output trees were visualized and beautified in the Interactive Tree Of Life (iTOL; v6)^70^.

### Calculation of evolutionary metrics

Filtered reads from each sample were mapped to all species-cluster representative MAGs concatenated together using Bowtie2 (v2.2.5; default parameters)^71^. Population statistics and nucleotide metrics including linkage disequlibrium (D’), nucleotide diversity (SNVs/kbp), nonsynonymous to synonymous mutation ratio (pN/pS) and major allele frequency were calculated from these mappings using the profile module of the inStrain program (v1.5.4; –database mode; default parameters)^15^ at genome and gene levels. Genetic annotation of MAGs was performed with Prodigal (v2.6.3; –p meta)^72^ for the gene module of inStrain.

### Inferring recombination rates

Filtered reads were mapped to each of 39 selected representative MAGs using Bowtie2 (v2.2.5; –sensitive-local mode)^71^. The mcorr package (https://github.com/kussell-lab/mcorr)^73^ was used to calculate the rate of recombination to mutation (gamma/mu) for each population. MAGs in which (1) normally distributed residuals for the model fit and (2) the bootstrapping mean was within 2X of the final estimate for gamma/mu^17^ were retained, resulting in a set of 10 genomes for inferring recombination rates.

### Statistical analyses

Statistical analysis was carried out in R (v4.0.0). Shapiro-Wilk and Bartlett’s tests were used to assess the normality and variance homogeneity of the data. The Kruskal-Wallis rank sum test with Chi-square correction was used for comparison of evolutionary metrics in genomes and genes among different groups. Pearson’s product-moment correlation was performed to assess the relationship between various evolutionary metrics (D’, pN/pS, r/m, coverage and SNVs/kbp for genomes; pN/pS, major allele frequency, coverage and SNVs/kbp for genes) and their relationship with sediment depth. Linear regression was used to fit the data and predict the linear correlation between the two indexes mentioned above on population. These metrics were used to test the evolutionary processes in the cold seep sediment populations and the effect of sediment depth on them.

## Data availability

MAGs, files for the phylogenetic trees and other related information have been uploaded to figshare (DOI: 10.6084/m9.figshare.17195003).

## Acknowledgements

The work was supported by the Scientific Research Foundation of Third Institute of Oceanography, MNR (No. 2022025), the National Natural Science Foundation of China (No. 41906076), the Science and Technology Projects in Guangzhou (No. 202102020970), and Guangdong Basic and Applied Basic Research Foundation (No. 20201910240000691).

## Author contributions

XD, ZS and CRJH designed this study. XD and YP performed analysis. XD, YP, MW, LW, WW, KJ and CG interpreted the data. YW, XX, JL and CRJH contributed to data collection. XD, YP and LW wrote the paper, with input from other authors.

