## Supplementary Figures for "Evolutionary ecology of microbial populations inhabiting deep sea sediments associated with cold seeps"

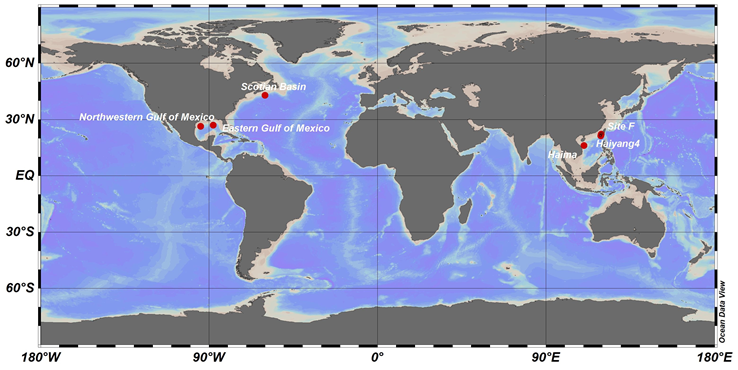

**Supplementary Figure 1. Geographical distributions of six cold seep sites analyzed in this study.** These sites were as follows: Eastern Gulf of Mexico; Northwestern Gulf of Mexico; Scotian Basin; Haiyang4, Site F, and Haima cold seeps in the South China Sea. Further details for each metagenome can be found in **Supplementary Table 1**.

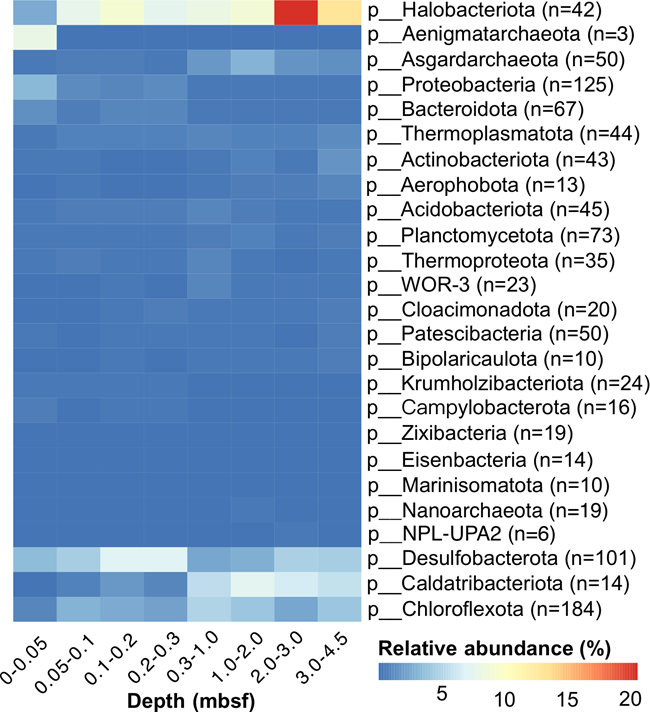

**Supplementary Figure 2. Relative abundances of cold seep sediment microbial communities at phylum level.** The 68 samples were catalogued into eight depth groups: 0-0.05 mbsf (n = 11); 0.05-0.1 mbsf (n = 14); 0.1-0.2 mbsf (n = 16); 0.2-0.3 mbsf (n = 8); 0.3-1.0 mbsf (n = 6); 1.0-2.0 mbsf (n = 4); 2.0-3.0 mbsf (n = 4); 3.0-4.5 mbsf (n = 5). The reported total relative abundance of each phylum was averaged from depth groups. Detailed data for relative abundance of each population in each sample can be found in **Supplementary Table 4**.

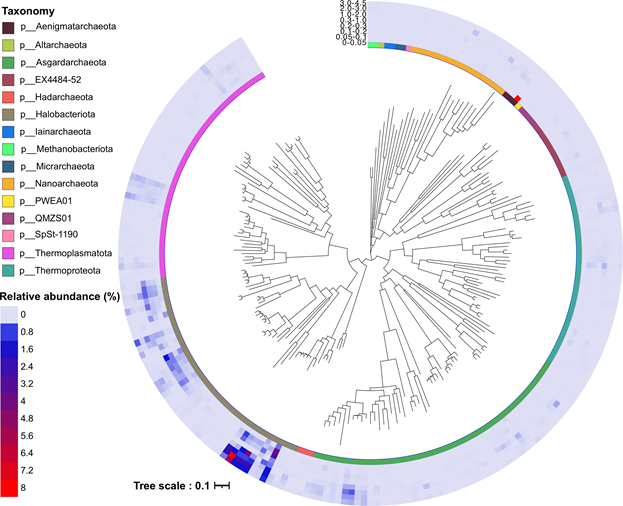

**Supplementary Figure 3. Maximum likelihood phylogenetic tree of assembled archaeal MAGs with their relative abundance.** The maximum-likelihood phylogenomic tree was built based on concatenated amino acid sequences of 122 archaeal marker genes produced by GTDB-Tk. Scale bar indicates the mean number of substitutions per site. See **Supplementary Figure 2** and **Supplementary Table 4** for more details about depth-profile relative abundances.

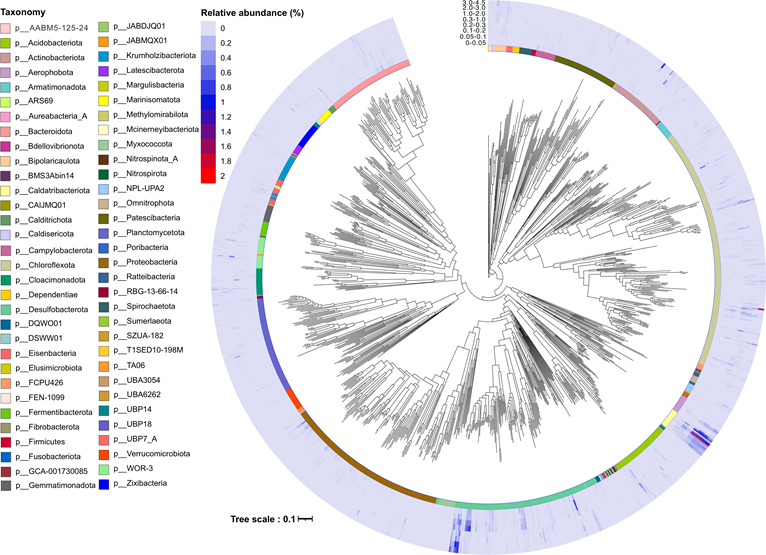

**Supplementary Figure 4. Maximum likelihood phylogenetic tree of assembled bacterial MAGs with relative abundance.** The maximum-likelihood phylogenomic tree was built based on concatenated amino acid sequences of 120 bacterial marker genes produced by GTDB-Tk. Scale bar indicates the mean number of substitutions per site. See **Supplementary Figure 2** and **Supplementary Table 4** for more details about depth-profile relative abundances.

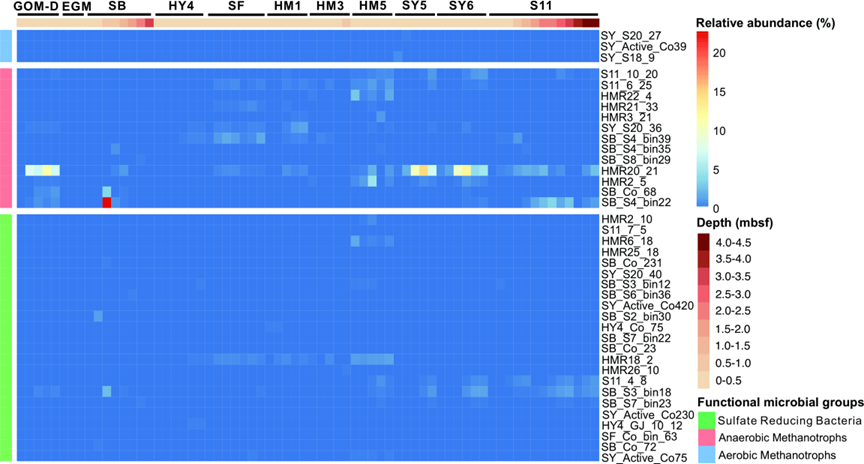

**Supplementary Figure 5. Relative abundances of 39 selected microbial species used for microdiversity analysis.** These species belong to aerobic methane-oxidizing bacteria, anaerobic methanotrophic archaea and sulfate-reducing bacteria, that are three key groups of functional microorganisms in cold seeps sediments. Detailed data for relative abundance of each population in each sample can be found in **Supplementary Table 4**.

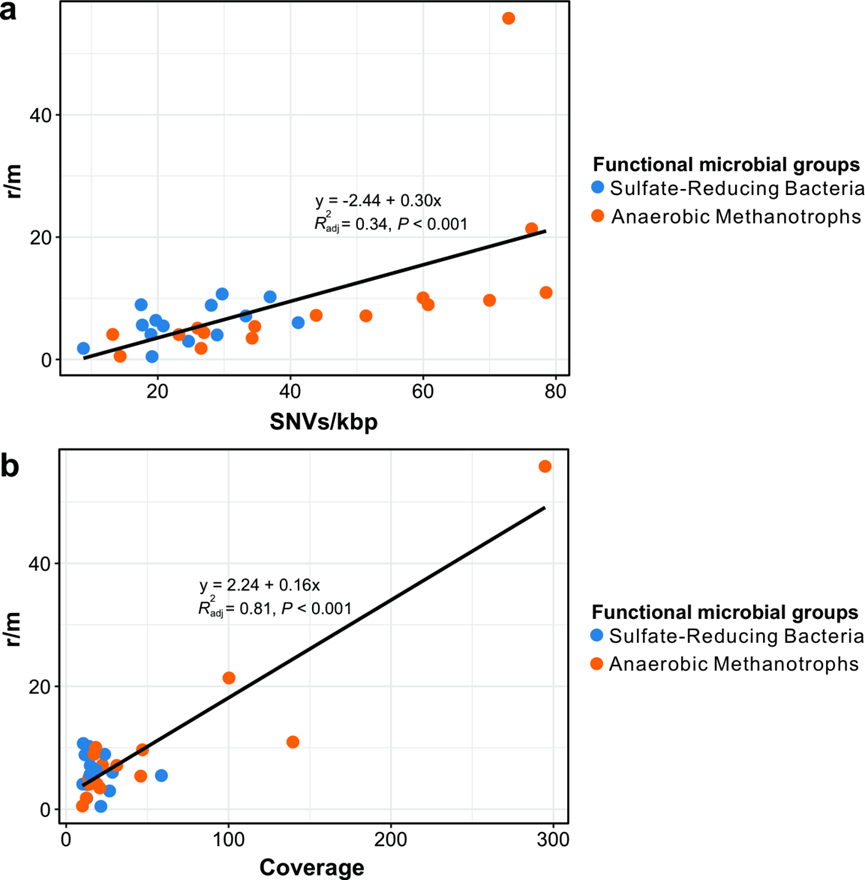

**Supplementary Figure 6. Genome-wide comparison of evolutionary metrics for anaerobic methanotrophic archaea and sulfate-reducing bacteria in cold seep sediments.** (a) Ratio of recombination to mutation (r/m) in relation to SNV density. (b) Ratio of r/m in relation to genome coverage. Each dot represents one species-level microbial population. Linear regressions and R^2^ values are indicated for different taxonomic groups. Detailed statistics for linear regressions are provided in **Supplementary Table 7**.

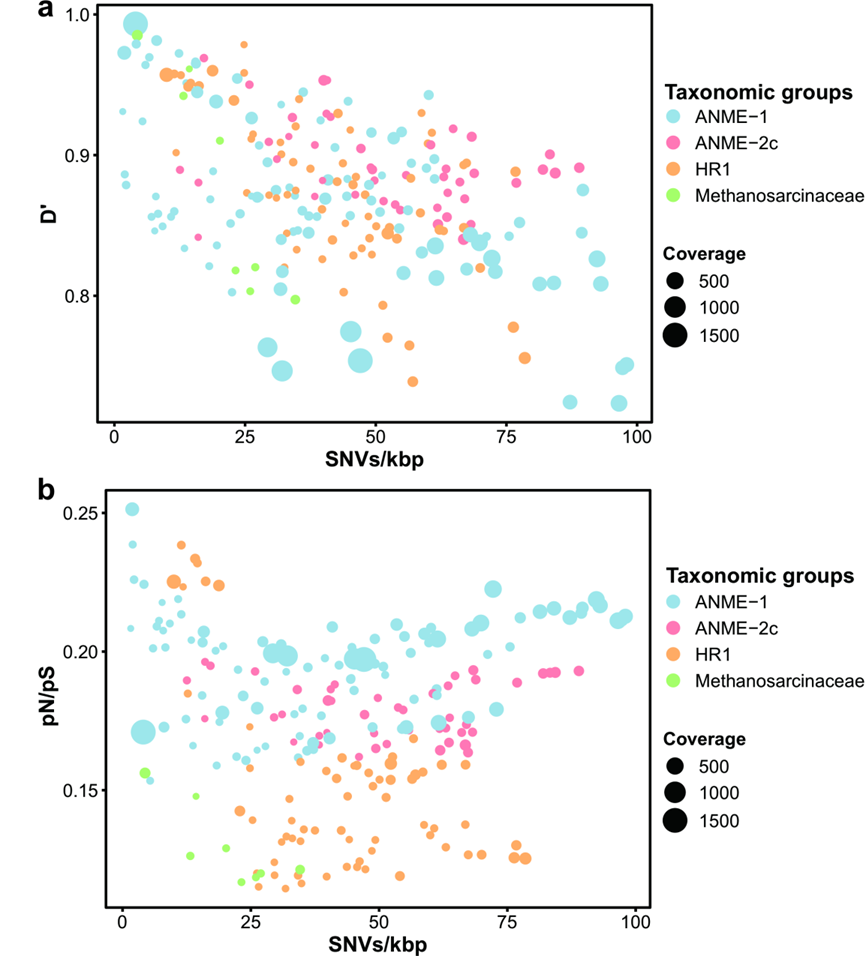

**Supplementary Figure 7. Genome-wide evolutionary metrics of anaerobic methanotrophic archaea in cold seep sediments.** (a) Relationships between SNV density, linkage disequilibrium (D’) and genome coverage. (b) Relationships between SNV density, pN/pS and genome coverage. Each dot represents one species-level microbial population. Source data can be found in **Supplementary Table 5**.

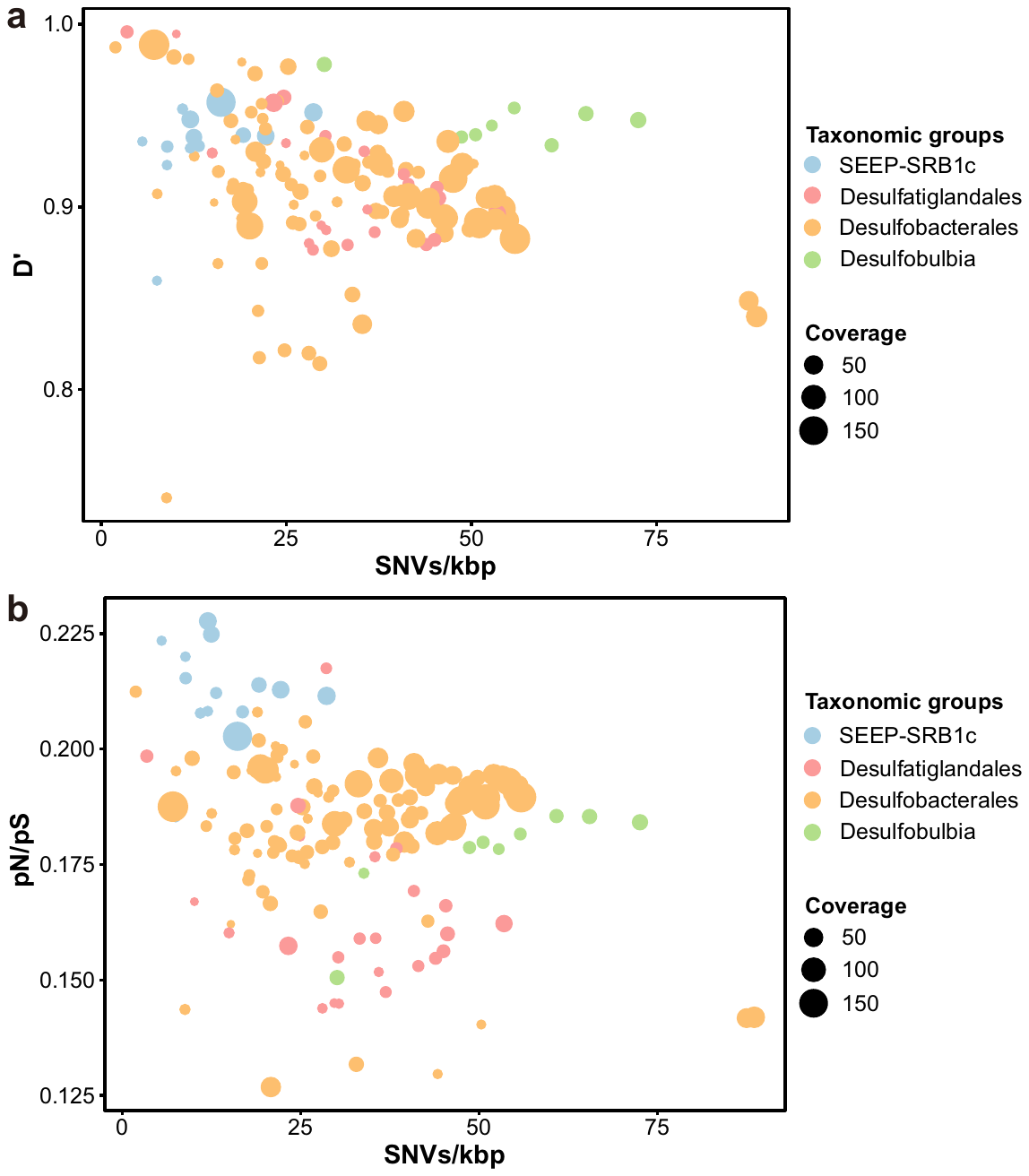
**Supplementary Figure 8. Genome-wide evolutionary metrics of sulfate-reducing bacteria in cold seep sediments.** (a) Relationships between SNV density, linkage disequilibrium (D’) and genome coverage. (b) Relationships between SNV density, pN/pS and genome coverage. Each dot represents one species-level microbial population. Source data can be found in **Supplementary Table 5**.

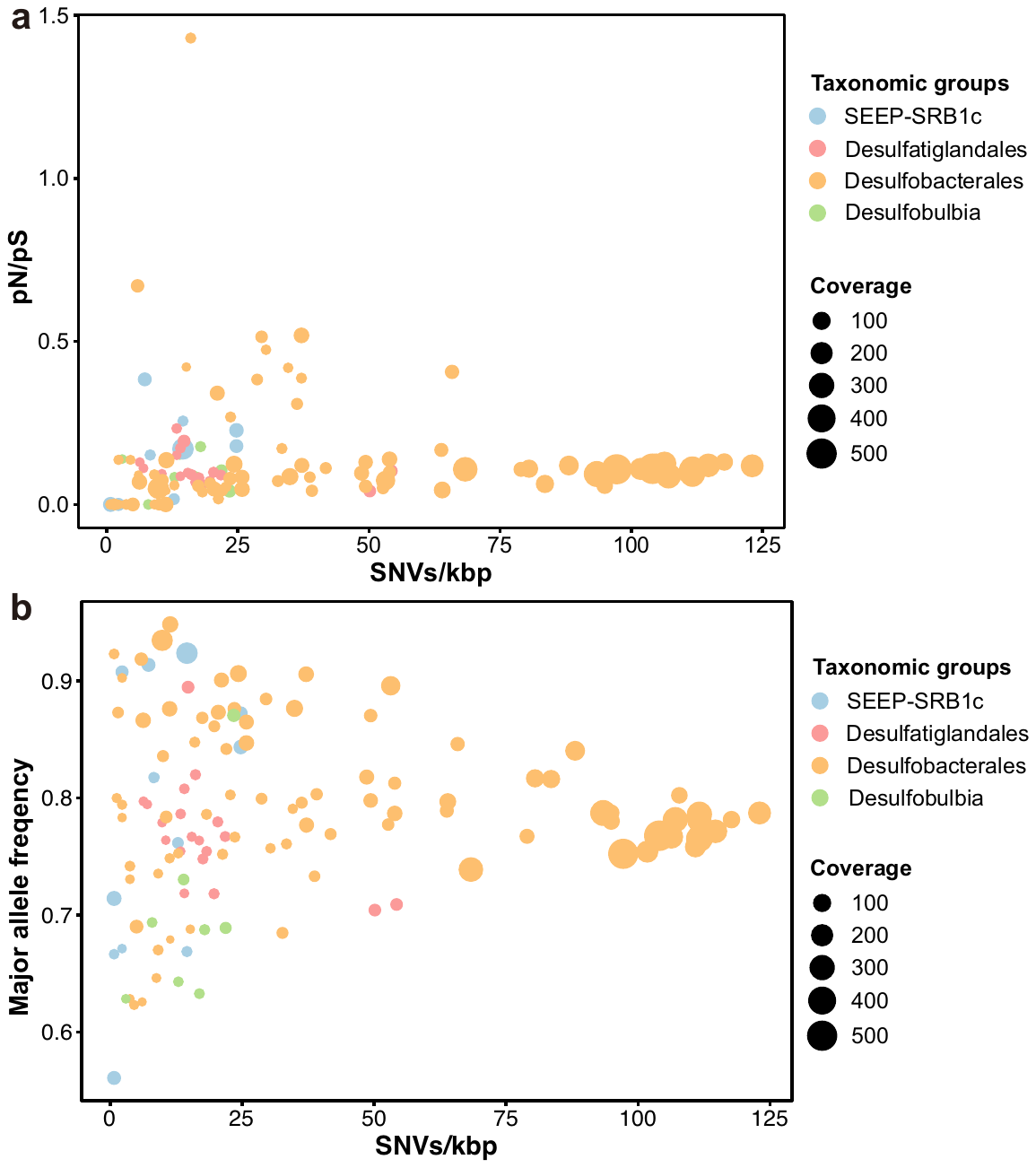
**Supplementary Figure 9. Gene-specific evolutionary metrics of sulfate-reducing bacteria in cold seep sediments.** (a) Relationships between SNV density, pN/pS and gene coverage at gene level. (b) Relationships between SNV density, major allele frequency and gene coverage at gene level. Each dot represents one species-level microbial population. Source data can be found in **Supplementary Table 8**.

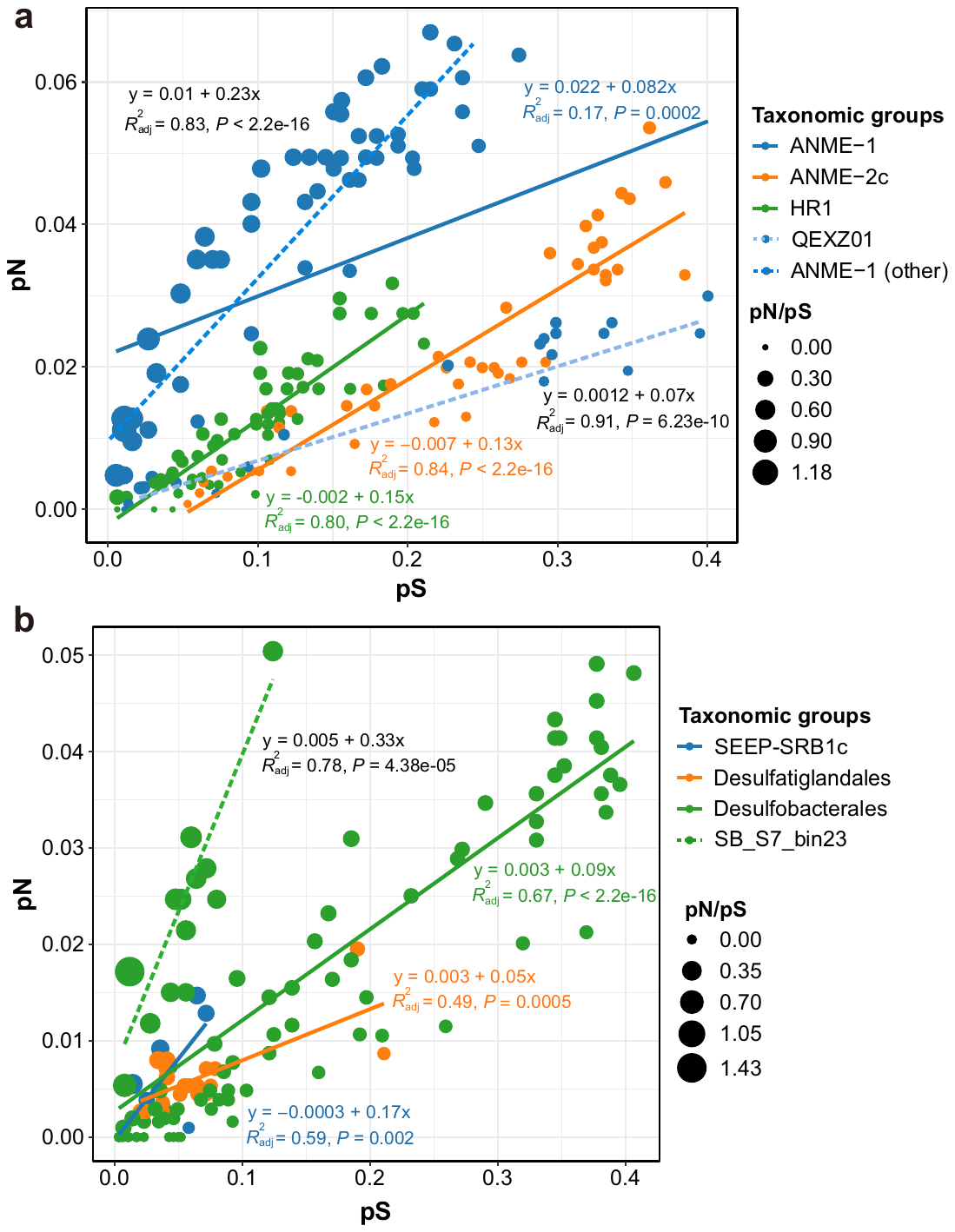
**Supplementary Figure 10. Comparison between pN and pS of functional genes in cold seep sediments.** (a) Linear regressions for *mcrA* genes from different taxonomic groups. (b) Linear regressions for *dsrA* genes from different taxonomic group. Detailed statistics for linear regressions are provided in **Supplementary Table 6**. Source data can be found in **Supplementary Table 8**.

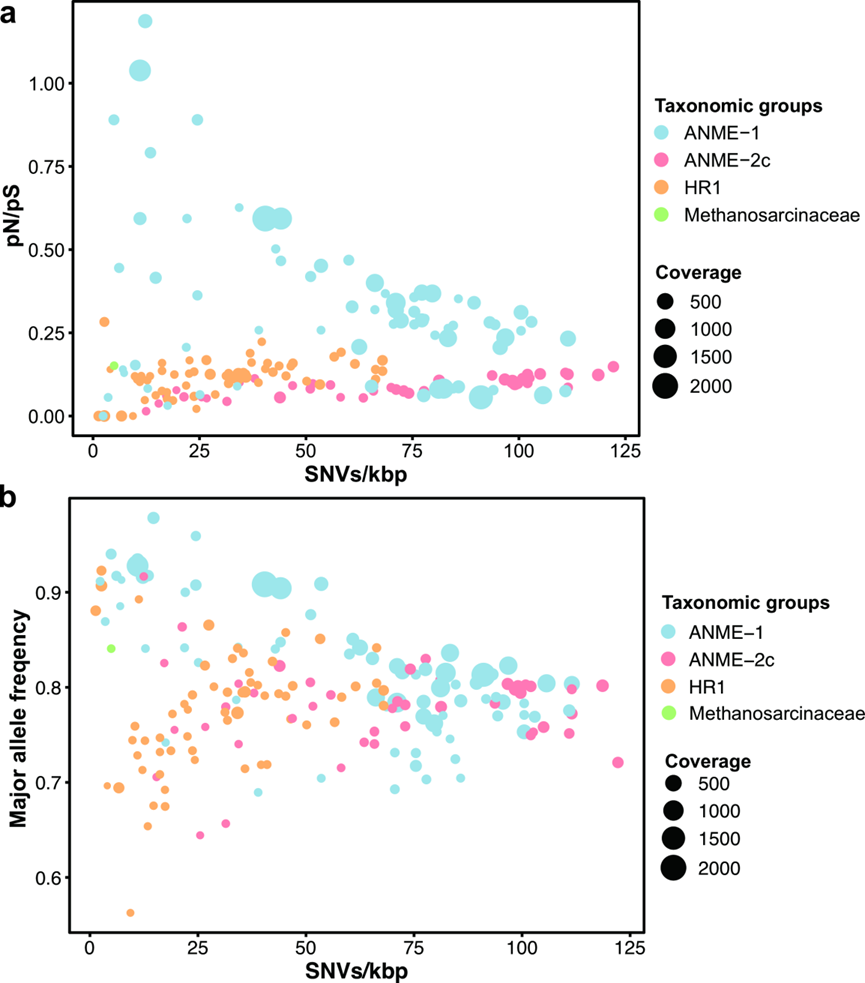

**Supplementary Figure 11. Gene-specific evolutionary metrics of anaerobic methanotrophic archaea in cold seep sediments.** (a) Relationships between SNV density, pN/pS and gene coverage at gene level. (b) Relationships between SNV density, major allele frequency and gene coverage at gene level. Each dot represents one species-level microbial population. Source data can be found in **Supplementary Table 8**.

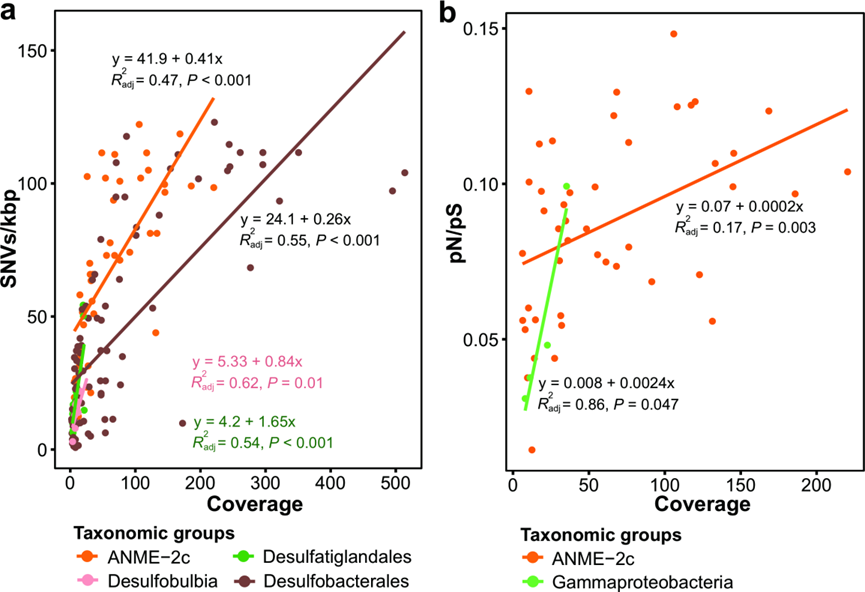

**Supplementary Figure 12. Comparison of evolutionary metrics for microbial populations in cold seep sediments at the gene level.** (a) SNV density in relation to gene coverage. (a) pN/pS in relation to gene coverage. Each dot represents one species-level microbial population. Linear regressions and R^2^ values are indicated for different taxonomic groups. Detailed statistics for linear regressions are provided in **Supplementary Table 6**.

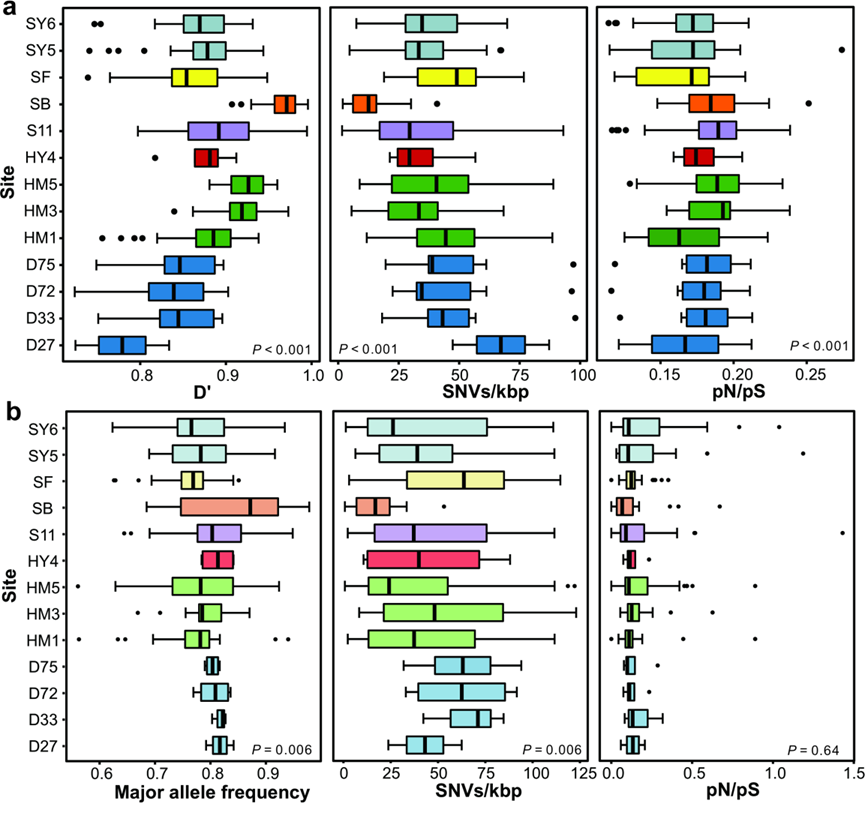

**Supplementary Figure 13. Site-specific comparison of evolutionary metrics for three key functional microbial groups in cold seep sediments.** (a) Box plot showing D’, SNV density and pN/pS ratio at genome level across different cold seep sites. (b) Box plot showing SNV density, pN/pS and major allele frequency of key functional genes (*pmoA*, *dsrA* and *mcrA*) across different cold seep sites. P-values of differences across different taxonomic groups were calculated using Kruskal-Wallis rank sum test. Source data can be found in **Supplementary Tables 5 and 8**. Detailed statistics for linear regressions are provided in **Supplementary Table 10**.
